# The Biofilm Lifestyle Shapes the Evolution of β-Lactamases

**DOI:** 10.1101/2023.10.02.560492

**Authors:** Øyvind M. Lorentzen, Anne Sofie B. Haukefer, Pål J. Johnsen, Christopher Frøhlich

## Abstract

The evolutionary relationship between the biofilm lifestyle and antibiotic resistance enzymes remains a subject of limited understanding. Here, we investigate how β-lactamases affect biofilm formation in *Vibrio cholerae* and how selection for a biofilm lifestyle impacts the evolution of these enzymes. Seven genetically diverse β-lactamases expressed in *V. cholerae* displayed a strong inhibitory effect on biofilm production, ranging from 17% to 61%. To understand how natural evolution affects this antagonistic pleiotropy under biofilm selecting conditions, we randomly mutagenized one β-lactamase and selected for elevated biofilm formation. Our results revealed that biofilm evolution selects for mutations predominantly clustered around the β-lactamase’s active site, yielding functional variants still proficient in β-lactam hydrolysis without biofilm inhibition. Mutational analysis of evolved variants demonstrated that restoration of biofilm development could be achieved either independent of enzymatic function or by actively leveraging enzymatic activity to increase biofilm formation. Taken together, the biofilm lifestyle can impose a profound selective pressure on antimicrobial resistance enzymes. Shedding light on such evolutionary interplays is of great importance to understand the various factors driving antimicrobial resistance.

**Impact statement:** β-lactamases inhibit biofilm formation and the selection for increased biofilm production can mitigate this antagonistic pleiotropic effect. The emergence of β-lactamase variants avoiding biofilm inhibition strongly suggests that the biofilm lifestyle affects the evolutionary fate of these enzymes.

## Introduction

Biofilms, structured bacterial communities covered in a protective extracellular matrix, represent one of the most prevalent bacterial lifestyles ^1,2^. Biofilm-embedded bacteria demonstrate a remarkable ability to endure harsh conditions and exhibit increased tolerance towards external stressors, including antimicrobials ^1,2^. These structured communities further serve as hotspots that facilitate the dissemination of mobile genetic elements harbouring antimicrobial resistance genes ^3–5^. It has been shown that a biofilm lifestyle can select for distinct evolutionary trajectories and profoundly influences the evolution of both bacteria and mobile genetic elements, when compared to bacteria evolving in unstructured environments ^5–8^. However, our current understanding of how biofilms influence the evolution of antimicrobial resistance enzymes is limited.

Upon acquisition, plasmid-harboured genes can induce pleiotropy, resulting in unpredictable effects on multiple cellular traits such as reduced basal bacterial growth or collateral responses to antimicrobials ^9–11^. Consequently, pleiotropy plays a pivotal role in shaping natural selection in a given environment, potentially requiring compensatory mutations to counteract these adverse effects. Among Gram-negative pathogens, the most prominent cause of β-lactam resistance is the production of β-lactamases ^12^. These enzymes display significant sequence- and functional-variability, and are often encoded on mobile genetic elements, which facilitates horizontal transmission to closely and more distantly related bacteria ^5^. They can be classified into Ambler classes A to D based on sequence diversity or grouped into two major functional categories: serine-type (classes A, C, and D) and metallo-β-lactamases (class B) ^13^. Enzymes classified under class A and D have been shown to antagonize biofilm formation in *Escherichia coli* and *Pseudomonas aeruginosa* ^14,15^. We hypothesize that the occurrence of such pleiotropic effects can significantly alter the evolutionary trajectory of the pleiotropy-inducing resistance enzymes.

In this study, we utilize *Vibrio cholerae* as a model organism, due to the significance of biofilm in its life cycle, to study the impact of β-lactamases on biofilm formation ^16–19^. We employed a combination of directed and experimental evolution techniques to evaluate how selection for pellicle production, a specific type of biofilm at the air-liquid interface, influences the evolutionary trajectories of β-lactamases (Fig. 1a) ^7,20,21^. Gaining insights into these intricate evolutionary relationships is essential for comprehending the dissemination and evolution of antimicrobial resistance enzymes.

**Fig. 1.**
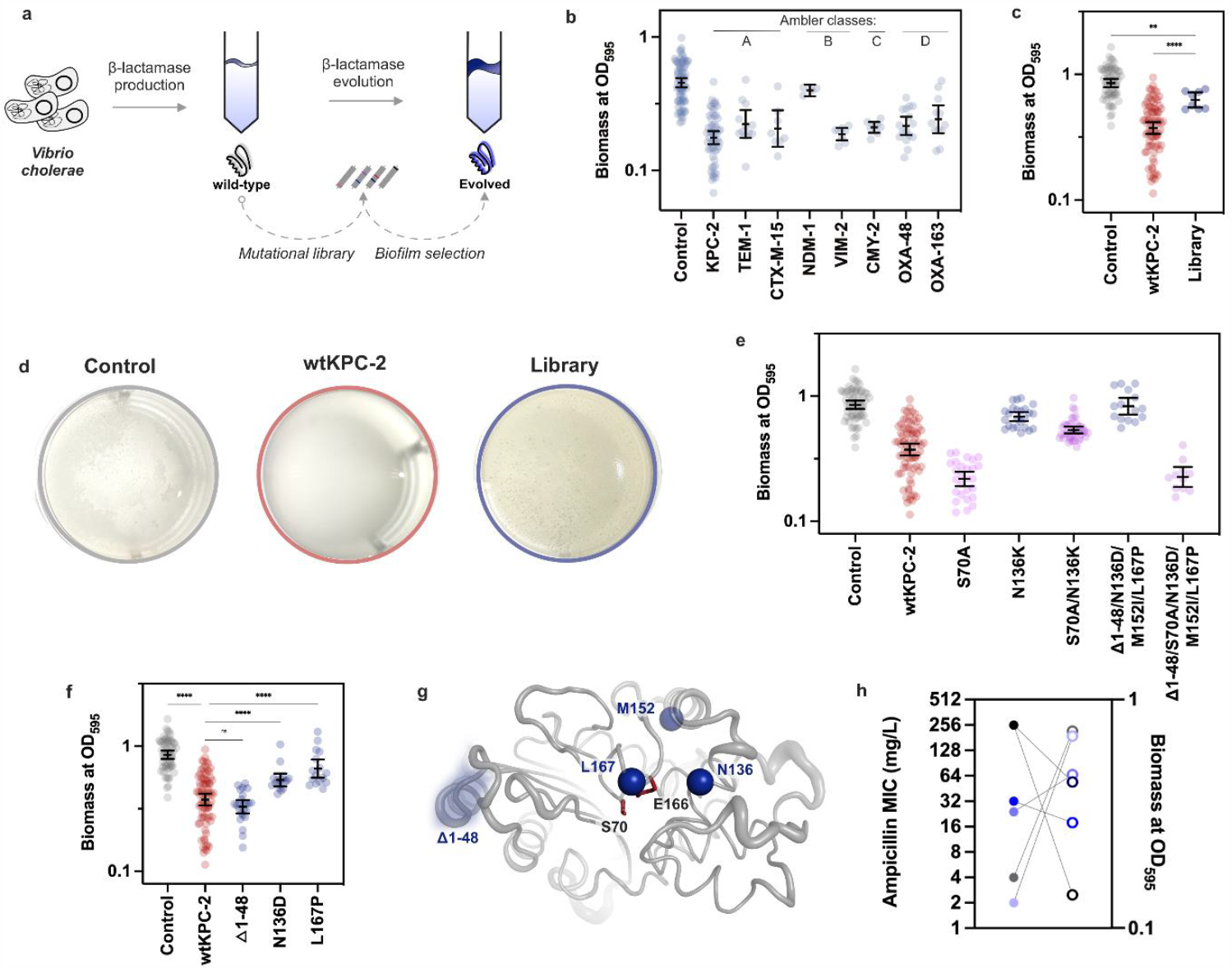
Biofilm lifestyle shapes the evolution of β-lactamases in *V. cholerae*. **a**. We first explored the influence of β-lactamase gene expression on the *V. cholerae* biofilm phenotype (left). Second, subjecting a mutant library of one β-lactamase to experimental evolution, we revealed how the biofilm lifestyle affects β-lactamase evolution (right). **b**. The expression of β-lactamase genes from Ambler classes A to D (top) significantly hindered biofilm formation in *V. cholerae* compared to the control vector **c**. Our mutational library of KPC (>5000 mutants) exhibited significantly enhanced biofilm formation compared to wtKPC-2 (****; one-way ANOVA, *P* < 0.0001), although it remained less than the vector control (**; one-way ANOVA, *P* = 0.001). **d**. Differences in biomass production related to *V. cholerae’s* ability to form pellicles. While our control displayed signs of pellicle formation after 24 h incubation, the presence of KPC-2 completely suppressed biofilm pellicle development. In contrast, the presence of our KPC-2 mutational library resulted in well-structured biofilm pellicles. **e**. While wtKPC-2 led to reduced biofilm capacity, N136K and Δ1-48/N136D/M152I/L167P, which were selected from and were enriched in the pellicle, demonstrated significant improvement in biofilm formation (*P* values reported in Tab. 2). β-lactam binding-deficient (serine-to-alanine at position 70) variants of wtKPC-2 and Δ1-48/N136D/M152I/L167P strongly reduced the biofilm phenotype. On the contrary, S70A/N136K maintained high levels of biofilm formation compared to the evolved variant N136K. **f**. Deconvolution of mutations within Δ1-48/N136D/M152I/L167P displayed that, in contrast to N136D and L167P, the deletion Δ1-48 did not significantly increase biofilm formation compared to wtKPC-2 (*P* values reported in Tab. 2). **g**. Location of mutational sites compared to the key active site residues S70 and E166 (red). **h**. Relationship between ampicillin resistance (filled circles) and biomass (empty circles) for wtKPC-2 (black), control (grey) and evolved mutants Δ1-48/N136D/M152I/L167P, N136K, N136D and L167P (displayed from light to dark blue, respectively). Each datapoint in b, c, e and f represents a biological replicate and error bars display 95% confidence intervals.

## Results and Discussion

### β-lactamases from all Ambler classes inhibit biofilm formation

To determine the inhibitory effect of β-lactamases on biofilm formation, we quantified biomass produced by *V. cholerae* strains harbouring a low-copy number vector with or without β-lactamase genes (Fig. 1a and b). Crystal violet staining of adherent biomass after 24 h of static growth was used as proxy for biofilm development. Compared to the control vector, seven out of eight tested β-lactamase-producing strains exhibited a significant reduction in biomass ranging from 43% to 61% (Tab. 1 and Fig. 1b, one-way ANOVA, *P* < 0.0001). Notably, NDM-1 was the only exception, showing a statistically non-significant reduction of 17% (Tab. 1 and Fig. 1b, one-way ANOVA, *P* > 0.05). While it has been previously suggested that biofilm inhibition is mainly attributed to class A and D β-lactamases due to their evolutionary relationship to low-molecular weight penicillin-binding proteins^14,15^, our data demonstrate that this antagonistic pleiotropy is much more general across the different classes of β-lactamases.

To exclude that the observed biofilm inhibition was due to intracellular protein aggregation or fitness defects, we determined ampicillin resistance and bacterial fitness of the β-lactamase-producing strains. Our data show that the β-lactamases conferred a 6- to 64-fold decrease in ampicillin susceptibility compared to our vector control, confirming the functional activity of the enzymes (Tab. 1). Evaluation of the area under the growth curve, used as a proxy for bacterial fitness, uncovered that seven out of eight β-lactamase-producing strains did not suffer a detrimental effect on fitness (Tab. 1, Fig. S1 and Tab. S1). In addition, there was no significant correlation between reduced bacterial fitness and biofilm formation (Pearson correlation, *R*^2^ = 0.15, *P* = 0.31, Fig. S1). Therefore, with TEM-1 as the exception, β-lactamase-mediated biofilm antagonization was likely not due to non-functional proteins (e.g., due to aggregation) or detrimental effects on bacterial fitness, but seemingly related to their enzymatic activity.

**Table 1.** Biofilm inhibitory effect of β-lactamases from all Ambler classes.

| Strain no. | β-lactamases | Ambler class | Biomass at $OD_{595}$<sup>a</sup> | *P* values<sup>b</sup> | *N*<sup>c</sup> | Relative fitness<sup>d</sup> | Ampicillin MICs (mg/L) |
| --- | --- | --- | --- | --- | --- | --- | --- |
| 30-71 | Control<sup>e</sup> | - | 0.454 ± 0.019 | - | 77 | 1 | 4 |
| 30-70 | KPC-2 | A | 0.176 ± 0.011 | <0.0001 | 56 | 1.34 ± 0.01 | >256 |
| 32-56 | TEM-1 | A | 0.223 ± 0.028 | 0.0002 | 12 | 0.80 ± 0.01 | 24 |
| 32-57 | CTX-M-15 | A | 0.206 ± 0.034 | 0.0033 | 8 | 1.22 ± 0.01 | >256 |
| 32-58 | NDM-1 | B | 0.398 ± 0.012 | 0.1264 | 4 | 1.07 ± 0.1 | >256 |
| 32-59 | VIM-2 | B | 0.187 ± 0.008 | <0.0001 | 8 | 1.26 ± 0.01 | >256 |
| 32-53 | CMY-2 | C | 0.210 ± 0.008 | <0.0001 | 8 | 0.94 ± 0.03 | 32 |
| 32-54 | OXA-48 | D | 0.216 ± 0.016 | <0.0001 | 16 | 1.32 ± 0.01 | >256 |
| 32-55 | OXA-163 | D | 0.242 ± 0.030 | 0.0007 | 12 | 0.97 ± 0.03 | >256 |

<sup>a</sup> *OD* measurements were performed in a 96-well plate.
<sup>b</sup> Brown-Forsythe one-way ANOVA to test for differences in biomass formation compared to the vector control ( $\alpha=0.05$ ) and followed by Dunnett test to correct for multiple testing; *F* (DFn, DFd) = 48.50 (8, 77); overall *P* value < 0.0001.
<sup>c</sup> Sample size (biological replicates) tested to determine the biomass at $OD_{595}$ .
<sup>d</sup> Determined as area under the growth curve compared to control (see Fig. S1 for bacterial growth curves and Tab. S1 for areas under curve).
<sup>e</sup> *V. cholerae* harbouring the pA15 vector without β-lactamase gene.
MIC = Minimal inhibitory concentration.
Errors are reported as the standard error of the mean.

### The biofilm lifestyle shapes the evolution of β-lactamases

To understand whether genetic changes within β-lactamases could modulate biofilm formation, we focused on the contemporary β-lactamase KPC-2 (wtKPC-2), as it conferred a strong biofilm inhibitory effect, and employed random mutagenizes as a means to generate genetic diversity (Fig. 1a). Expression of the gene library in *V. cholerae* significantly improved biofilm formation, resulting in higher biomass (*OD*_595_) compared to the wtKPC-2 (Fig. 1c, one-way ANOVA, *P* < 0.0001). Improvements were also evident in the strains’ ability to form biofilm pellicles (Fig. 1d). The presence of wtKPC-2 completely suppressed pellicle formation, while both the control and mutant library formed visible biofilm pellicles at the air liquid interface. Thus, our results indicate that the library harbours *bla*_KPC-2_ mutants able to compensate for the initial biofilm antagonization.

To identify potential variants displaying compensatory behaviour, we harvested the biofilm pellicles formed by the library population-mix (n=2) and isolated random clones. Sanger sequencing revealed that 33% of the selected clones contained mutations within KPC-2 at position 136 (N136K and N136D), indicating strong selection for pellicle production and parallel evolution. The clone displaying N136D also exhibited additional amino acid substitutions (M152I/L167P) and a deletion which led to a frameshift mutation (Fig. S2). This frameshift resulted in the loss of the first 48 amino acids, including the signal peptide, and recruitment of an alternative methionine start codon at position 49 (Δ1-48/N136D/M152I/L167P).

To remove the effect of potential confounding mutations on the vector backbone or chromosome, we sub-cloned N136K and Δ1-48/N136D/M152I/L167P into an isogenic vector backbone and assayed biofilm formation. N136K and Δ1-48/N136D/M152I/L167P displayed 68% and 107% higher biomass relative to wtKPC-2, respectively (Tab. 2 and Fig. 1e). To deconvolute the contributions from the different mutations in the Δ1-48/N136D/M152I/L167P variant, we selected and constructed mutants displaying either loss of the signal peptide (Δ1-48) or mutations around the active site (N136D and L167P; Tab. 2 and Fig. 1f). While Δ1-48 alone did not significantly (one-way ANOVA, *P* = 0.79) improve biofilm formation relative to wtKPC-2, N136D and L167P increased biofilm formation by 33% (*P* = 0.0003) and 67% (*P* < 0.0001), respectively (Tab. 2 and Fig. 1f). Translocation of β-lactamases into the periplasmic space depends on the presence of a signal peptide and the loss of a signal peptide (e.g., Δ1-48/N136D/M152I/L167P) prevents this. Thus, KPC-2 variants without this signal peptide are likely retained in the cytoplasm. Our data demonstrate that these mutants still exert a detrimental effect on biofilm formation. In addition, β-lactamases bearing a signal peptide (e.g., N136D/K and L167P) can be functionally folded and enzymatically active within the cytosol before the translocation ^22^. This observation suggests that the mechanistic interaction by which they interfere with biofilm formation likely occurs in the cytoplasm. Taken together, our result show that a biofilm lifestyle can select for mutations in β-lactamases that reverse their initial antagonistic pleiotropic effect on biofilm formation. Thus, evolution in biofilms shapes the evolution of antibiotic resistance enzyme.

**Table 2.** KPC-2 variants resulting in improved biofilm development.

| Strain no. | Inserts | Biomass at $OD_{595}$<sup>a</sup> | *P* values<sup>b</sup> | *N*<sup>c</sup> | Relative fitness<sup>d</sup> | Ampicillin MICs (mg/L) |
| --- | --- | --- | --- | --- | --- | --- |
| 30-70 | wtKPC-2 | 0.417 ± 0.020 | - | 81 | 1 | >256 |
| 30-71 | Control<sup>e</sup> | 0.889 ± 0.032 | <0.0001 | 61 | 0.75 ± 0.01 | 4 |
| 30-65 | S70A | 0.229 ± 0.014 | <0.0001 | 27 | ND | 4 |
| 30-77 | Δ1-48/N136D/M152I/L167P<sup>e</sup> | 0.862 ± 0.063 | <0.0001 | 15 | 0.78 ± 0.04 | 2 |
| 30-67 | Δ1-48/S70A/N136D/M152I/L167P<sup>e</sup> | 0.235 ± 0.021 | 0.0008 | 11 | ND | 4 |
| 32-13 | Δ1-48 | 0.341 ± 0.019 | 0.7914 | 24 | 0.79 ± 0.03 | 2 |
| 32-07 | N136D | 0.553 ± 0.037 | 0.0003 | 16 | 0.97 ± 0.02 | 32 |
| 30-75 | N136K | 0.698 ± 0.028 | <0.0001 | 24 | 1.00 ± 0.01 | 24 |
| 30-69 | S70A/N136K | 0.547 ± 0.019 | <0.0001 | 39 | ND | 2 |
| 32-06 | L167P | 0.696 ± 0.061 | <0.0001 | 16 | 0.99 ± 0.01 | >256 |

<sup>a</sup> *OD* measurement was performed in 24-well plates.
<sup>b</sup> Brown-Forsythe one way ANOVA to test for differences in biomass formation compared to wtKPC-2 ( $\alpha=0.05$ ) and followed by Dunnett test to correct for multiple testing; *F* (DFn, DFd) = 62 (15, 293); overall *P* value < 0.0001.
<sup>c</sup> Sample size (biological replicates) tested to determine the biomass at $OD_{595}$ .
<sup>d</sup> Determined as area under the growth curve compared to wtKPC-2 (see Fig. S3 for bacterial growth curves and Tab. S1 for area under curves)
<sup>e</sup> *V. cholerae* harbouring the pA15 vector without β-lactamase gene
MIC = Minimal inhibitory concentration.
ND = Not determined
Errors are reported as the standard error of the mean.

### Functional mutations in KPC-2 reverse biofilm inhibition

Next, we investigated the functional and structural role of mutations acquired during evolution. To study the functionality of selected and constructed mutants, we investigated their ability to confer ampicillin resistance (Tab. 2). As expected, all mutants lacking the signal peptide exhibited antibiotic resistance similar to the vector control. Other mutations, such as N136D/K and L167P, clustered around the active site of KPC-2 (Fig. 1g) maintained ampicillin resistance 6- to >64-fold higher than the control strain, demonstrating that biofilm compensation can occur without complete loss of their enzymatic activity (Fig. 1h) ^23,24^. Furthermore, the fitness effect of these mutants did not significantly correlate with biofilm formation (Pearson correlation, *R*^2^ =0.004 and *P* =0.89, Fig. S3). Altogether, our combined findings on bacterial growth and biofilm formation indicate that reversal of biofilm inhibition was not related to changes in bacterial fitness.

Our mutant analysis (Fig. 1e) indicated that the compensatory effects of the evolved mutants may be linked to cytosolic processes, where they could either be permanently active due to loss of the signal peptide (Δ1-48/N136D/M152I/L167P) or temporarily (N136D/K and L167P) prior to translocation ^22^. To investigate whether the enzymatic activity of wtKPC-2 and evolved variants was linked to biofilm formation, we constructed serine-to-alanine mutants at position 70 which are unable to covalently bind and efficiently hydrolyze β-lactam substrates (Tab. 2 and Fig. 1e) ^25^. As expected, introducing S70A in the wtKPC-2, N136K, and Δ1-48/N136D/M152I/L167P backgrounds resulted in MICs similar to the vector control (Tab. 2). Introducing S70A in wtKPC-2 induced a further reduction of 45% in biofilm formation compared to the wtKPC-2. Similarly, the introduction of S70A in the evolved Δ1-48/N136D/M152I/L167P variant caused a 73% decrease in biofilm formation compared to the evolved variant, resulting in biofilm levels similar to KPC-2:S70A (Fig. 1e). On the contrary, S70A/N136K displayed only a 21% reduction in biofilm formation compared to the evolved N136K variant and maintained a strong biofilm phenotype relative to the other serine-to-alanine mutants. Our findings stand in contrast to a previous study where the removal of the active site serine either rescued (TEM-1) or had no effect (OXA-3) on biofilm formation ^14^. Consequently, how enzyme functionality affects biofilm inhibition seems to vary greatly between different β-lactamases. While the exact molecular mechanisms of biofilm antagonization and compensation remain elusive, informed by our data we hypothesize that the evolved variants compensate biofilm inhibition either independent of enzymatic activity (N136K) or by actively leveraging its enzymatic activity (Δ1-48/N136D/M152I/L167P). For the latter, the active site serine seems to be recruited during evolution and is indispensable for restoration of biofilm formation (Fig. 1e).

Taken together, our findings demonstrate that a broad range of highly diverse β-lactamases inhibits biofilm formation in *V. cholerae*, and that selection for a biofilm lifestyle significantly affects the evolution of these enzymes. Such pleiotropy, where genes can affect a multitude of bacterial phenotypes, has been observed in multiple model systems ^26–28^. We argue that the selection pressure generated through pleiotropic effects represents a substantial selective force, which influences the genetic adaption and evolution of antimicrobial resistance enzymes.

## METHODS AND MATERIAL

### Growth media and chemicals

All strains and primers used and constructed within this study are shown in supplementary Tab. S2 and S3. Strains were grown in Lysogeny-Broth (LB) media supplemented with chloramphenicol (5 or 25 mg/L for *V. cholerae* and *E. coli* strains, respectively). LB media and chloramphenicol were purchased from Sigma-Aldrich (USA). Restriction enzymes and T4 ligase were supplied by ThermoFisher (USA).

### Strain construction

The gene sequences of *bla*_CMY-2_, *bla*_CTX-M-15_ and *bla*_NDM-1_ were previously synthesized by Genewiz (Germany) and subcloned in a low copy number vector (pA15 origin) according to the gene sequences NG_048935.1, NG_048814.1 and NG_049326.1, respectively ^29^. In addition, *bla*_OXA-48_ (CP033880) and *bla*_KPC-2_ (KU665642) were earlier subcloned from *E. coli* 50579417 and *Klebsiella pneumoniae* K47-25, respectively, into the same vector backbone ^29–31^. All diverse β-lactamases were modified and carried an additional glycine after their start codon allowing us to use a *Xho*I restriction site at the N-terminus.

We subcloned *bla*_VIM-2_ (NG_050347.1) from the *Pseudomonas aeruginosa* K34*-7* using P105/P106 ^32^. *bla*_TEM-1_ was synthesized (Genewiz) according to the gene sequence NG_050145.1 and amplified using P90/P91 ^32^. Amplification was performed with Phusion polymerase (NEB). PCR products were digested using *Dpn*I, *Xho*I, and *Nco*I and ligated with backbone using T4 Ligase. Ligated vectors were transformation into the *E. coli* E. cloni (MP-21-5) and subsequently transformed into *V. cholerae* C6706.

OXA-163 was constructed by site directed mutagenesis and whole vector amplification using Phusion polymerase (NEB, USA), primers P54F/R and *bla*_OXA-48_(CP033880) as a template ^33^. PCR product was digested for 1 h at 37°C using *Dpn*I and *Lgu*I. The digested product was ligated for 1 h at room temperature using T4 ligase and transformed into *E*.*coli* E.cloni. Cells were selected on chloramphenicol 25 mg/L and target gene Sanger sequenced.

To subclone mutant *bla*_KPC-2_ alleles, the target genes and vector backbone were amplified using primers P7/P8 and P3/P4, respectively (Tab. S3), and Phusion polymerase (NEB). PCR products were digested using *Dpn*I, *Xho*I, and *Nco*I and ligated with backbone using T4 Ligase. Ligated vectors were transformation into the *E. coli* E. cloni (MP-21-5) and subsequently transferred into *V. cholerae* C6706.

Active-site serine of KPC-2 was mutated to alanine (S70A) using whole vector site-directed mutagenesis with primers P108/P115 containing *Lgu*I cutting sites to investigate whether disruption of the enzymatic activity would affect biofilm formation. The *bla*_KPC-2_ genes were amplified using primers P108/P115 and Phusion polymerase (NEB). The PCR products were digested with *Lgu*I and *Dpn*I for 1 hour at 37°C following self-ligation using T4 ligase. Ligated vectors were transformation into MP-21-5 and subsequently transferred into *V. cholerae* C6706.

### Bacterial fitness measurements

Single colonies were grown in overnight under shaking at 700 rpm and 37 °C and subsequently diluted 1:100 into LB-medium supplemented with 5mg/L chloramphenicol to a final volume of 300 μL. Growth curve experiments were conducted in Bioscreen 100-well-honeycomb plate (Oy Growth Curves Ab Ltd, Finland) where the OD_600_ was monitored in four-minute intervals using a Bioscreen plate reader (Oy Growth Curves Ab Ltd). Growth curves were recorded over 18 h at 37°C with continuous shaking. The relative bacterial fitness was calculated as area under the curve of the individual growth curves using the flux package in R ^34^ and normalized to either *V. cholerae* harbouring an empty control vector (Tab. 1) or wtKPC-2 (Tab. 2). Fitness was calculated based on a minimum of three biological replicates each determined based on three technical replicates per biological replicate.

### Biomass determining using crystal violet

Overnight cultures were prepared in 2 mL LB medium supplemented with 5 mg/L chloramphenicol and incubated overnight at 37°C with shaking (700 rpm). The following day, the cultures were diluted 1:100 in 2 mL LB medium in a 24-well plate (Corning, USA) and incubated statically at 37°C for 24 h. Pellicle formation was imaged with a NexiusZoom stereo microscope (Euromex, Netherlands) at 6.7x magnification. Next, the bacterial cultures were removed from the 24-well plate and the plate was gently washed in distilled water to remove non-adherent bacterial cells. Biofilms were fixed by incubation for 1 h at 55°C. To quantify the attached biomass, cells were stained with 2 mL of 0.1% crystal violet (Sigma-Aldrich) for 10 min. The crystal violet solution was removed, and the plates were washed in filtered water and airdried. Crystal violet-stained biomass was dissolved in 2.25 ml 70% ethanol (Sigma-Aldrich) and quantified by determining the OD_595_ in a Spark^®^ multimode plate reader (Tecan, Switzerland). Datasets were tested for normality using a Shapiro-Wilk test (α=0.05). The log transformed datasets were analyzed using a Brown-Forsythe one-way analysis of variance (ANOVA, α=0.05) followed by a Dunnett test to corrected for multiple comparisons tests. All statistical analyses were performed using Prism v. 9 (GraphPad, USA).

### MIC determination

To assess the functionality of the constructed β-lactamases in *V. cholerae* (Tab. 1 and 2), antimicrobial susceptibility against ampicillin was determined using MIC Test Strips (Liofilchem, Italy). Briefly, a bacterial suspension with an optical density of 0.5 McFarland (1.5x 10^8^ CFU/mL) was prepared in 0.9% saline (Sigma-Aldrich). In order to ensure maintenance of the pA15 vector, LB agar plates supplemented with 5 mg/L chloramphenicol and were inoculated with the prepared bacterial suspensions. The ampicillin MIC was visually determined after incubation for 20 h at 37°C.

### Mutagenesis and biofilm selection

The KPC mutant library used in this study was constructed using error-prone PCR to introduce mutations in the *bla*_KPC-2_ gene as previously described ^29^. Briefly, Mutational libraries were constructed by error-prone PCR using 10 ng vector DNA, GoTag DNA polymerase (Promega, USA), 25 mM MgCl_2_ (Promega), 10 μM of primers P7/P8 and either 50 μM oxo-dGTP or 1 μM dPTP. PCR products were *Dpn*I digested for 1 h at 37°C. 5 ng of each product was used for a second PCR, which was performed as described above, but without mutagenic nucleotides. PCR product from the 2^nd^ PCR was digested using *Nco*I and *Xh*oI and ligated in a 1:3 ratio with the digested and purified vector backbone. The resulting ligation mixture was transformed into MP21-5 (*E. coli* E. cloni® 10G). To ensure that the entire sequence space could be sampled, >5000 mutants were harvested. DNA from the mutational library was isolated and 10 ng was electroporated into *V. cholerae* C6706 (resulting in MP30-72), and selected on LB plates containing 5 mg/L chloramphenicol (>5000 colonies).

Overnight cultures of MP30-72 were prepared in 3 mL LB medium (n=2) supplemented with 5 mg/L chloramphenicol and incubated overnight at 37°C with shaking (700 rpm). Cultures were diluted 1:100 in 2 mL LB and chloramphenicol in a 24-well plate and incubated statically at 37°C for 48 h. To harvest biofilm pellicles, a sterile inoculation loop was used to transfer biofilm pellicles into 1 mL phosphate saline buffer (Fisher Bioreagents, USA, 0.137 M NaCl, 0.0027 M KCl and 0.01 M phosphate, pH 7.4). Afterwards, the suspension was homogenised for 120 s to dislodge biofilm-embedded bacterial cells. To isolate single biofilm-evolved clones, 1 μL of the bacterial suspension was spread onto LB agar supplemented with 5 mg/L chloramphenicol and incubated overnight at 37°C. Single clones were randomly harvested and changes in the target gene identified using Sanger sequencing (Genewiz). KPC-2 genes carrying mutations were subcloned into isogenic backgrounds and their ability to form biomass was determined, all as described above.

## Funding and Acknowledgment

The authors declare no conflict of interest. Funding was obtained from the Centre for new antibacterial strategies (CANS) at UiT - The Arctic University of Norway (UiT) (CF, ASBH). ØML was funded by UiT. PJJ thanks The Olav Thon Foundation for funding. We thank Rebekka Rolfsnes for assisting in strain construction.

