## Supplementary information for "The Biofilm Lifestyle Shapes the Evolution of β-Lactamases"

### Contents

1. Supplementary tables 2
   Table S1: Area under the growth curves 2
   Table S2: Strains constructed and used in this study. 3
   Table S3: Primers used in this study 4
2. Supplementary figures 5
   Figure S1: Fitness effect of β-lactamases in *V. cholerea*. 5
   Figure S2: Multiple sequence alignment of wtKPC-2 and evolved variants 6
   Figure S3: Fitness effect of KPC-2 variants in V. cholerea. 7
3. References 8

### 1. Supplementary tables

**Table S1: Area under the growth curves**

| Strain no. | Insert | Area under the growth curve[a,b] | *N* |
|---|---|---|---|
| 30-70 | wtKPC-2 | 1251 ± 6.1 | 14 |
| 30-71 | Control | 932.8 ± 12.1 | 6 |
| 32-56 | TEM-1 | 745.4 ± 6.6 | 6 |
| 32-57 | CTX-M-15 | 1136 ± 11.2 | 6 |
| 32-58 | NDM-1 | 1001 ± 95 | 6 |
| 32-59 | VIM-2 | 1175 ± 4.2 | 6 |
| 32-53 | CMY-2 | 878.9 ± 28.0 | 6 |
| 32-54 | OXA-48 | 1229 ± 12.9 | 6 |
| 32-55 | OXA-163 | 919.6 ± 36.3 | 6 |
| 30-77 | Δ1-48/N136D/M152I/L167P | 975 ± 46.3 | 5 |
| 32-13 | Δ1-48 | 988.5 ± 40.6 | 3 |
| 32-07 | N136D | 1211 ± 26.6 | 3 |
| 30-75 | N136K | 1256 ± 4.2 | 8 |
| 32-06 | L167P | 1243 ± 5.0 | 3 |

[a] Growth curves are determined in *V. cholerae* C6706
[b] Error is given as the standard error of the mean.

**Table S2: Strains constructed and used in this study.**

| Strain no. | Strain background [a] | Vector number | Insert | Reference |
|---|---|---|---|---|
| 30-73 | C6706 | None | None | [1] |
| 30-71 | C6706 | pUNS-233 | pA15 vector without *bla* gene | This study |
| 30-70 | C6706 | pUNS-146.2 | KPC-2 | This study |
| 32-56 | C6706 | pUNS-239 | TEM-1 | This study |
| 32-57 | C6706 | pUNS-156 | CTX-M-15 | This study |
| 32-58 | C6706 | pUNS-157 | NDM-1 | This study |
| 32-59 | C6706 | pUNS-236 | VIM-2 | This study |
| 32-53 | C6706 | pUNS-158 | CMY-2 | This study |
| 32-54 | C6706 | pUNe-4 | OXA-48 | This study |
| 32-55 | C6706 | pUNS-178 | OXA-163 | This study |
| 30-72 | C6706 | pUNS-146.2 | Mutational library of pUN-*bla*$_{KPC-2}$ | This study |
| 30-65 | C6706 | pUNS-240 | KPC-2: S70A | This study |
| 30-76 | C6706 | pUNS-241 | KPC-2: Δ1-48/N136D/M152I/L167P | This study |
| 30-77 | C6706 | pUNS-241.1[b] | KPC-2: Δ1-48/N136D/M152I/L167P | This study |
| 30-67 | C6706 | pUNS-242 | KPC-2: Δ1-48/S70A/N136D/M152I/L167P | This study |
| 30-74 | C6706 | pUNS-238 | KPC-2: N136K | This study |
| 30-75 | C6706 | pUNS-238.1[b] | KPC-2: N136K | This study |
| 30-69 | C6706 | pUNS-243 | KPC-2: S70A/N136K | This study |
| 32-13 | C6706 | pUNS-245 | KPC-2: Δ1-48 | This study |
| 32-06 | C6706 | pUNS-247 | KPC-2: L167P | This study |
| 32-07 | C6706 | pUNS-248 | KPC-2: N136D | This study |
| | | **Cloning strains:** | | |
| 21-05 | *E. coli* E. cloni® 10G | None | None | Lucigen |
| 24-44 | *E. coli* E. cloni® 10G | pUNS-146.2 | KPC-2 (KU665642; modified) | [2] |
| 30-60 | *E. coli* E. cloni® 10G | pUNS-239 | TEM-1 (NG_050145.1) | This study |
| 24-80 | *E. coli* E. cloni® 10G | pUNS-156 | CTX-M-15 (NG_048814.1) | [2] |
| 24-81 | *E. coli* E. cloni® 10G | pUNS-157 | NDM-1 (NG_049326.1) | [2] |
| 30-57 | *E. coli* E. cloni® 10G | pUNS-236 | VIM-2 (NG_050347.1) | This study |
| 12-69 | *E. coli* E. cloni® 10G | pUNS-158 | CMY-2 (NG_048935.1) | [2] |
| 21-01 | *E. coli* E. cloni® 10G | pUNe-4 | OXA-48 (CP033880) | [2] |
| 29-27 | *E. coli* E. cloni® 10G | pUNS-178 | OXA-163 (CP033880; modified) | This study |
| 24-45 | *E. coli* E. cloni® 10G | - | KPC-2 mutational library | [2] |
| 30-64 | *E. coli* E. cloni® 10G | pUNS-240 | KPC-2: S70A | This study |
| 30-81 | *E. coli* E. cloni® 10G | pUNS-245 | KPC-2: Δ1-48 | This study |
| 30-66 | *E. coli* E. cloni® 10G | pUNS-242 | KPC-2: Δ1-48/S70A/N136D/M152I/L167P[c] | This study |
| 32-03 | *E. coli* E. cloni® 10G | pUNS-247 | KPC-2: L167P | This study |
| 32-23 | *E. coli* E. cloni® 10G | pUNS-238.1 | KPC-2: N136K | This study |
| 30-69 | *E. coli* E. cloni® 10G | pUNS-247 | KPC-2: S70A/N136K | This study |
| 32-04 | *E. coli* E. cloni® 10G | pUNS-248 | KPC-2: N136D | This study |
| 30-81 | *E. coli* E. cloni® 10G | pUNS-245 | KPC-2: Δ1-48 | This study |
| | | **Clinical isolates for strain constructions:** | | |
| K34-7 | *Pseudomonas* | - | Carrier of pVIM-2 | [3] |

[a] *V. cholerae* C6706 strain originates from El Tor biotype Inaba

[b] Target gene was subcloned after selection into an isogenic pA15 vector backbone and isogenic *V. cholerae* C6706 strain

**Table S3: Primers used in this study**

| No. | Name | | Sequence (5’ to 3’) | Ref. |
|---|---|---|---|---|
| 3 | pUN-NcoI | F | GCTTTCCCATGGATGTTTTTCCTCCTTATGTTAAGCTTACTCAG | [2] |
| 4 | pUN-XhoI | R | GCTTCTCGAGAAGTGGTTAGCGCGTATTTGTG | |
| 7 | preOXAseq | F | GATTACGCGCAGACCAAAACG | [2] |
| 8 | postOXAseq | R | CCTATTTCCCTAAAGGGTTTATTGAGAATATG | |
| 105 | NcoI-VIM-2 | F | TTTTTTGGCCATGGGATTCAAACTTTTGAGTAAGTTATTGGTCTATTTGACC | This study |
| 106 | XhoI-VIM-2 | R | TTTTTCTCGAGCTACTCAACGACTGAGCGATTTGTGTG | This study |
| 115 | KPC-2_S70A_LguI | F | TTTTTGCTCTTCTGTGCGCGTCATCAAGGGCTTTCTTGC | This study |
| 108 | KPC_2_S70A_LguI | R | TTTTTGCTCTTCGCACAGTGGGAAGCGCTCC | This study |
| 117 | KPC-ORF2-NcoI | F | GCTTCCATGGGAGATACCGGCTCAGGCGCAAC | This study |
| | | R | Primer number 4 | [2] |
| 118 | KPC-N136D-LguI | F | TTTTTGCTCTTCGCCGCCGCCGATTTGTTGCTGAAGG | This study |
| 118 | KPC-N136D-LguI | R | TTTTTGCTCTTCGCGGCGTTATCACTGTATTGC | This study |
| 119 | KPC-L167P-LguI | F | TTTTTGCTCTTCCTGGGAGCCGGAGCTGAACTCC | This study |
| 119 | KPC-L167P-LguI | R | TTTTTGCTCTTCCCCAGCGGTCCAGACGG | This study |
| 90 | NcoI-TEM-1 | F | TTTTTTCCATGGGAAGTATTCAACATTTTCGTGT | This study |
| 91 | XhoI-TEM-1 | R | TTTTTTCTCGAGTTACCAATGCTTAATCAGTG | This study |
| 54 | OXA-163-LguI- | F | TTTTTGCTCTTCTATTCGGGCTAAAACTGGATACGATACTAAGATTGGCTGG | This study |
| 54 | OXA-163-LguI | R | TTTTTGCTCTTCGAATAATATAGTCGCCATTG | This study |

### 2. Supplementary figures

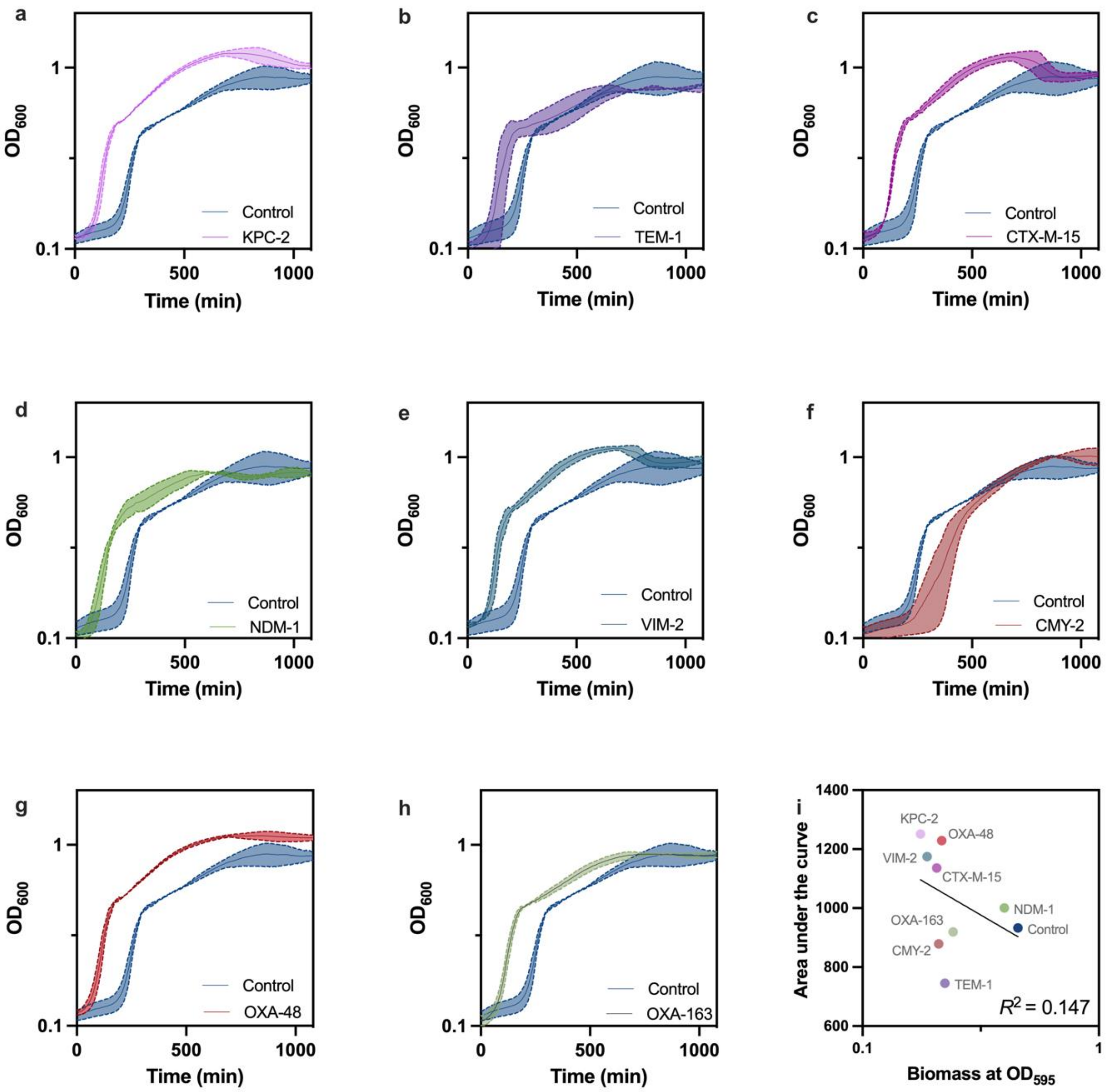


**Figure S1: Fitness effect of β-lactamases in *V. cholerea*.**

Effect of class A: KPC-2 **(a)**, TEM-1 **(b)**, CTX-M-15 **(c)**, class B: NDM-1 **(d)**, VIM-2 **(e)**, class C: CMY-2 **(f)** and class D: OXA-48 **(g)**, OXA-163 **(h)** β-lactamase and the empty vector control (n = 6) on bacterial fitness in *V. cholerae*. Bacterial fitness was assessed as the area under the growth curve over 18 h of incubation at 37°C. **i.** Non-significant correlation between area under the curves and the strains ability to form biomass (see Tab. 1) resulted in non-significant Pearson correlation ($R^2$ = 0.147, $P$ = 0.31). Error around the growth curves represents the standard error of the mean determined by at least 3 biological replicates. See table S1 for overview of $N$ for each strain.

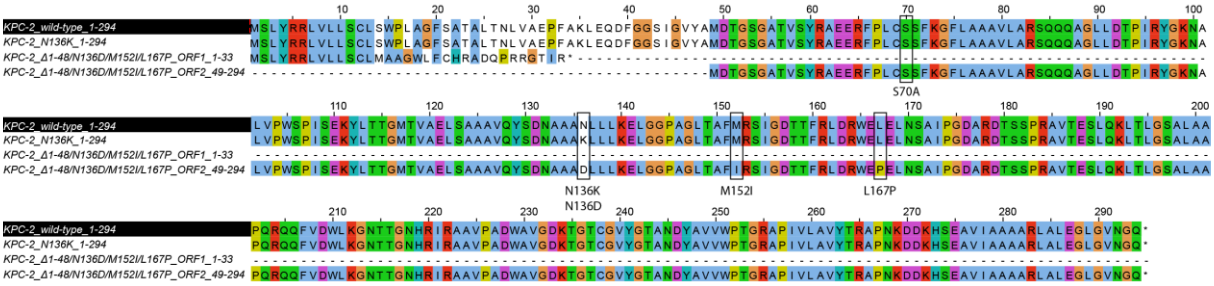


**Figure S2: Multiple sequence alignment of wtKPC-2 and evolved variants**

Both reading frames of Δ1-48/N136D/M152I/L167P are presented; the first reading frame (Δ1-48/N136D/M152I/L167P_ORF1) includes amino acids 1-33. The nucleotide deletions in nucleotide positions 40 and 41 (amino acid position 14) led to a frameshift which subsequently introduced a premature stop codon. The second reading frame (Δ1-48/N136D/M152I/L167P_ORF2) is introduced by a new start codon at position 49 and includes amino acids 49-294. Amino acid substitutions found in Δ1-48/N136D/M152I/L167P (N136D/M152I/L167P) and N136K (N136K), and the active site S70, are shown in brackets at their respective positions. The coloured residues are indicated as follows; Blue: hydrophobic. Red: positive charge. Magenta: negative charge. Green: polar. Orange: glycines. Yellow: prolines. Cyan: aromatic. White: not conserved.

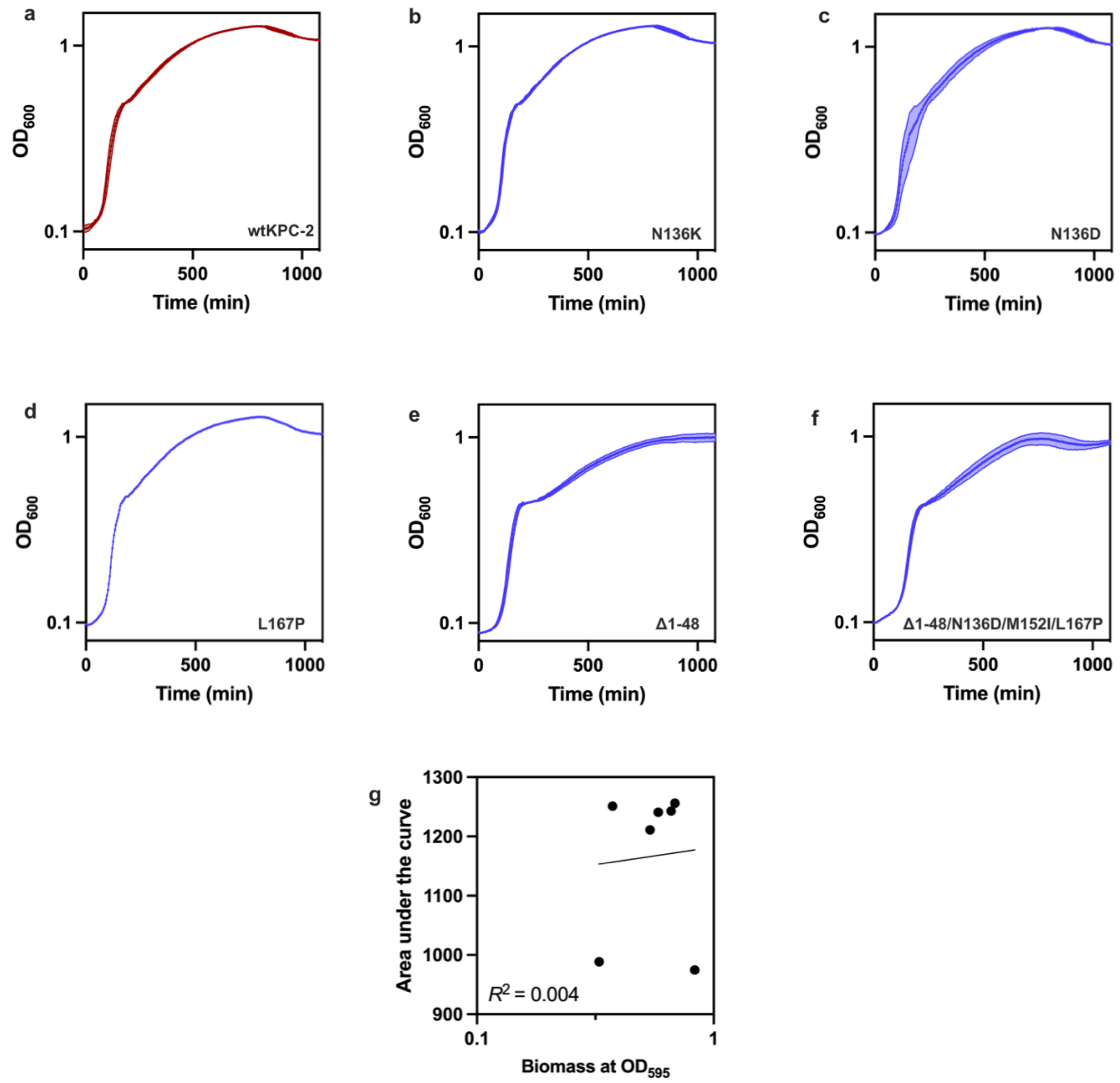


**Figure S3: Fitness effect of KPC-2 variants in V. cholerea.**

Growth curves of different KPC-2 mutants in V. cholerae. **a**. wild-type KPC-2, **b**. N136K, **c**. N136D, **d**. L167P, **e**. Δ1-48, **f**. Δ1-48/N136D/M152I/L167P and **g**. the non-significant correlation between area under the curve and the strains' biofilm biomass (see Tab. 2) (Pearson correlation, $R^2$ = 0.04, $P$ = 0.89). Error around the growth curves represents the standard error of the mean determined by at least 3 biological replicates. See table S1 for overview of *N* for each strain.
